# The clinical antiprotozoal drug halofuginone promotes weight loss by elevating GDF15 and FGF21

**DOI:** 10.1101/2024.08.18.608423

**Authors:** Suowen Xu, Zhenghong Liu, Tian Tian, Wenqi Zhao, Zhihua Wang, Monan Liu, Mengyun Xu, Fanshun Zhang, Zhidan Zhang, Meijie Chen, Yanjun Yin, Meiming Su, Wenhao Pan, Shiyong Liu, Min-dian Li, Peter J. Little, Danielle Kamato, Songyang Zhang, Dongdong Wang, Stefan Offermanns, John R. Speakman, Jianping Weng

## Abstract

Obesity is a debilitating disease with increasing worldwide prevalence. Despite its high prevalence, specific pharmacologic intervention for obesity is challenging. Here, we report that halofuginone, an FDA-approved anti-scleroderma and anti-protozoal drug, is a promising anti-obesity agent in rodent models. Halofuginone suppressed food intake, increased energy expenditure, and resulted in weight loss in preclinical diet-induced obese mouse models, while also decreasing insulin resistance and hepatic steatosis. By combining genetic and pharmacological tools with transcriptomics, we identified that halofuginone increases FGF21 and GDF15 levels via ATF4. Using knockout mice, we show these hormones are both necessary for its anti-obesity effects. Thus, our study first reports the beneficial metabolic effects of halofuginone and underscores its utility to treat obesity and its associated metabolic complications.

## Main Text

Obesity, characterized by the pathological and ectopic accumulation of adipose tissue due to relative overnutrition, is widely acknowledged as a chronic degenerative condition that predisposes individuals to various metabolic-related disorders, including cardiovascular disease, type 2 diabetes mellitus, metabolic dysfunction-associated steatotic liver disease and cancer ^1–3^. Obesity constitutes one of the greatest healthcare challenges and imposes a heavy economic burden on public health systems because of its rising prevalence ^4,5^. Efforts to manage obesity through lifestyle interventions and modifications, including dietary and physical activity changes, and early therapeutic interventions have achieved limited success. Bariatric surgery is successful but impractical to perform in the large population sectors that suffer from obesity, and has a high risk of side effects. Consequently, the development of new therapeutic agents and strategies is deemed a more effective approach for sustained obesity management at the population level ^6,7^. This is exemplified by the recent success of glucagon-like peptide-1 receptor (GLP1R) agonists^8,9^. But in a significant portion of the treated population, these drugs cause undesirable side effects^10^ underscoring the need to continuously discover and develop innovative drugs acting via novel pathways.

Restriction of dietary protein or certain amino acids, such as methionine and branched-chain amino acids (BCAAs), promotes metabolic health through pathways including GCN2/ATF4, mTOR, and AMPK^11–13^. Halofuginone (HF) is an FDA-approved drug used to treat scleroderma and protozoal infections. HF functions by inhibiting aminoacyl-tRNA synthetases, with a primary action on glutamyl-prolyl-tRNA synthetase (EPRS)^14^. This inhibition leads to activation of the amino acid starvation (AAS) response pathway and integrated stress response (ISR) signaling. Inhibition of EPRS by HF leads to the accumulation of uncharged tRNAs, initiating autophosphorylation of the amino acid sensor GCN2, resulting in phosphorylation of eIF2α and increased expression of ATF4. Beyond its anti-malarial properties, HF demonstrates a broad therapeutic potential against various medical conditions by its anti-inflammatory, anti-fibrotic, and anti-tumor effects ^15–18^. However, it remains unclear whether HF can exert metabolic benefits by treating obesity-associated metabolic disorders.

In this study, HF were identified as a repurposed drug for treating obesity by elevating GDF15 and FGF21 via ATF4 pathway, which respectively inhibit energy intake and increase energy expenditure, thus promoting weight loss and an improved metabolic profile in obese mouse models. Our findings opened up a hitherto unappreciated therapeutic avenue to alleviate obesity and associated metabolic disorders by the application of anti-protozoal drug halofuginone.

## Results

### Halofuginone promotes weight loss in obese animal models

Taking into consideration relevant preclinical and clinical dosing regimens^19,20^, we investigated the tissue distribution and pharmacokinetics of HF in mice. Following administration via intravenous (168 μg/kg) and oral (840 μg/kg) routes, we measured the concentrations of HF in various organs and blood compartments. After *i.v.* injection, HF was detected in the heart plasma, portal plasma, heart, liver, spleen, lung, kidney, fat and muscle. The drug concentration reached the highest at 15 min post injection (Fig. 1A). When given orally, HF mainly accumulated in the stomach and intestines (Fig. 1B). The oral bioavailability of HF was 60.4%. Two hours after oral administration, HF reached its peak plasma concentration (12.5 ± 2.9 ng/ml). The half-life of orally and intravenously administered HF was around 5.0 and 3.4 hours, respectively (Fig. 1C).

**Fig. 1.**
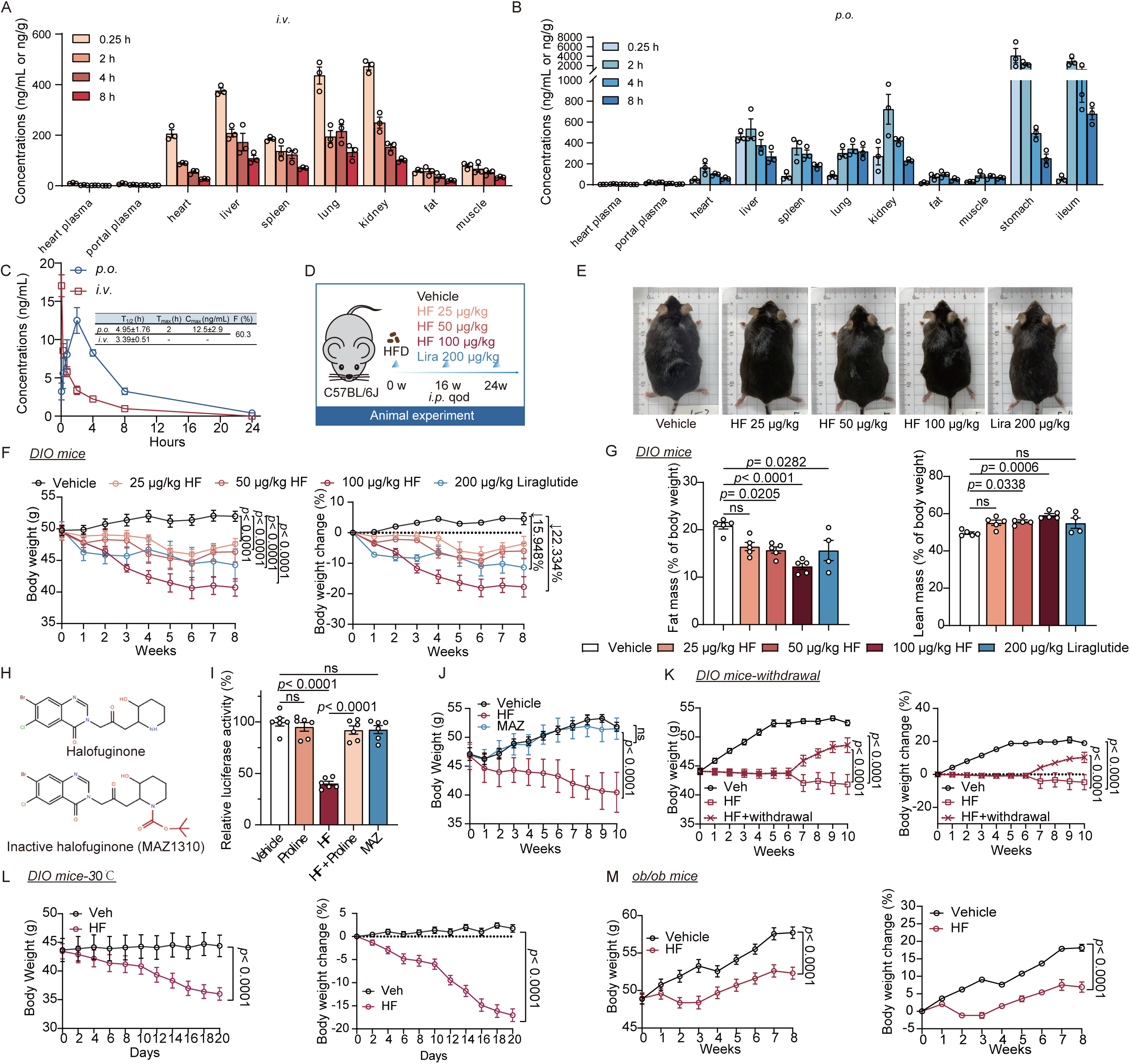
Halofuginone promotes weight loss in obese animal models. **(A)** Concentrations (ng/mL or ng/g) of HF in CD-1 mouse plasma and tissues after *i.v.* administration of HF (168 μg/kg, n=3). **(B)** Concentrations (ng/mL or ng/g) of HF in CD-1 mouse plasma and tissues after *p.o.* administration of HF (840 μg/kg, n=3). **(C)** Pharmacokinetics parameters of HF after *p.o* and *i.v.* administration in CD-1 mice (n=3). **(D-G)** 8-week-old male C57BL/6J mice were fed a HFD for 16 weeks, and then randomly assigned to four experimental groups: (1) vehicle group, (2) 25 μg/kg HF treatment group, (3) 50 μg/kg HF treatment group or (4) 100 μg/kg HF treatment group, (5) 200 μg/kg liraglutide treatment group. During HFD challenge, mice were treated with intraperitoneal injections of vehicle or indicated drugs every two days. (**D)** Schematic diagram illustrating DIO mice treated with escalating doses of HF or GLP1R agonist liraglutide. **(E)** Representative gross images of DIO mice after vehicle-, HF- or liraglutide-treatment. The mice were subjected to the following measurements: (**F)** Body weight and % body weight change (vehicle or HF group: n = 5, liraglutide group: n = 4). (**G)** Fat mass and lean mass (% of body weight) by NMR scans (vehicle or HF group: n = 5, liraglutide group: n = 4). **(H)** Chemical structure of HF and MAZ1310 (EPRS1 inactive compound). **(I)** Luciferase mRNA was incubated with rabbit reticulocyte lysate (RRL), and the translation process was measured using a luminescence assay (HF: 200 nM, proline: 8 mM, MAZ1310: 200 nM). **(J)** Body weight of DIO mice during vehicle, 100 μg/kg HF treatment or 100 μg/kg MAZ treatment (vehicle or MAZ group: n = 5, HF group: n = 4). **(K)** Body weight and % body weight change of DIO mice during vehicle, HF treatment or HF withdrawal treatment (n = 10). **(L)** Body weight and % body weight change of DIO mice in a thermoneutral environment (30LJ) during 20-days HF treatment (100 μg/kg, N=8). **(M)** Body weight and % body weight change of DIO mice during vehicle or HF treatment (vehicle group: n = 7, HF group: n=8, 100 μg/kg). Data in (G) and (I) were analyzed by one-way ANOVA followed by Bonferroni’s multiple-comparison test. Data in (F) and (J-M) were determined through two-way ANOVA followed by Bonferroni’s multiple-comparison test.

In clinical trials, for chronic administration of HF, oral dosing at 0.5 mg/day is recommended^19,21^. After dosage conversion between humans and mice^22^, we selected 100 μg/kg for further evaluation of the safety profile of HF in normal chow diet fed C57BL/6J mice (Fig. S1A). We observed comparable long-term weight gain between mice receiving 100 μg/kg HF via intraperitoneal injection every 2 days and the vehicle group (Fig. S1B). A subsequent comprehensive comparison of blood cell counts, histopathology, organ weights, and serum parameters indicative of injury in the liver, heart and kidney function between HF and vehicle suggested no noticeable toxicity of HF in mice (Fig. S1C-1R).

To determine whether HF exhibits anti-obesity effects, we administered escalating doses (25, 50, 100 μg/kg) of HF and 200 μg/kg of liraglutide (reference drug, a GLP1 receptor agonist) by intraperitoneal injection every two days for 8 weeks in diet-induced obese (DIO) mice (Fig. 1D-E). In the highest dosage group of HF treatment (100 μg/kg), the body weight of the HF treatment group (40.7 g, SD=2.8) decreased by 22.3% compared to the vehicle group (52.0 g, SD=1.6), whereas the positive control drug, liraglutide, resulted in a 15.9% reduction in body weight (44.3 g, SD=4.43) in mice (Fig. 1F). The 25 μg/kg and 50 μg/kg HF-treated groups also exhibited significant weight-lowering effects. Moreover, HF-treated mice exhibited a reduction in total fat mass without a corresponding impact on lean mass percentage, paralleling the outcomes observed in the liraglutide-treated group (Fig. 1G). Consistently, 100 μg/kg HF significantly reduced serum levels of triglycerides (TG) and total cholesterol (TC) (Fig. S2A-B). In addition, obese mice treated with either HF or liraglutide exhibited reduced levels of alanine aminotransferase (ALT) and aspartate aminotransferase (AST) (Fig. S2C-D). There was no significant effect of HF on creatinine (CREA) and UREA levels (Fig. S2E-F). Moreover, HF treatment was associated with a decrease in HOMA-IR and improvements in glucose tolerance and insulin sensitivity (Fig. S2G-J). We next asked whether or not EPRS inhibition by HF is essential for the weight loss effects of HF, by intraperitoneal injection of DIO mice with MAZ1310, an inactive HF derivative (unable to bind EPRS) (Fig. 1H)^23^. Through *in vitro* translation experiments using rabbit reticulocyte lysate, we demonstrated that HF can inhibit translation, and this effect can be reversed by the addition of excess proline (Fig. 1I). We used MAZ1310 as a negative control in mouse dosing experiments. Compared to an equivalent dose of HF (100 μg/kg), MAZ1310 did not cause body weight loss (Fig. 1J).

HF-treated mice regain some of their lost body weight after discontinuation of the treatment (Fig. 1K). The mice in the HF-withdrawal group have less body weight (compared with vehicle) but gained about 10% body weight over a period of four weeks. The benefits of HF on reducing body weight also pertain to female DIO mice, which exhibited reduced body weight and enhanced insulin sensitivity (Fig. S2K-L). In the oral HF experiment, the weight loss achieved with HF at 100 μg/kg (-9.8%) was comparable to that observed with orlistat at 20 mg/kg (-8.2%). However, HF at 200 μg/kg (-27.9%) demonstrated a significantly enhanced weight loss effect (Fig. S2M). HF also reduced the rate of weight gain in DIO mice housing under thermoneutral (TN) conditions (-18.8%) and *ob/ob* mice (-11.25%) (Fig. 1L-M).

### Halofuginone suppresses food intake and increases energy expenditure

HF (100 μg/kg) administration inhibited cumulative food intake, as well as food intake in fast-refeeding experiment (Fig.2A-B). The decline in the body weight was greater following HF treatment than that in pair-fed in mice (Fig.2C). This suggested the impact of HF was due to a combination of inhibited food intake and elevated energy expenditure. Moreover, HF treatment led to improvements in thermogenic capability, as shown by sustained elevated core temperature during cold challenge (Fig.2D). Of note, western blot analysis showed that UCP1 was upregulated in iBAT of HF-treated obese mice versus vehicle (Fig.2E). Energy expenditure of DIO mice treated with vehicle or HF also supported the findings that HF elevated energy expenditure (Fig.2F). HF exhibited significant impacts on food intake and prompted the utilization of fat as the primary energy source as revealed by RER analysis (Fig.2G). However, in mice fed normal chow diet, there was no significant effect on energy expenditure (p > .05), despite the fact that food intake was notably suppressed (Fig. S3A-C). Collectively, these findings demonstrate that HF significantly ameliorates obesity-related phenotypes, suppresses food intake, and enhances energy expenditure in obese mice.

**Fig. 2.**
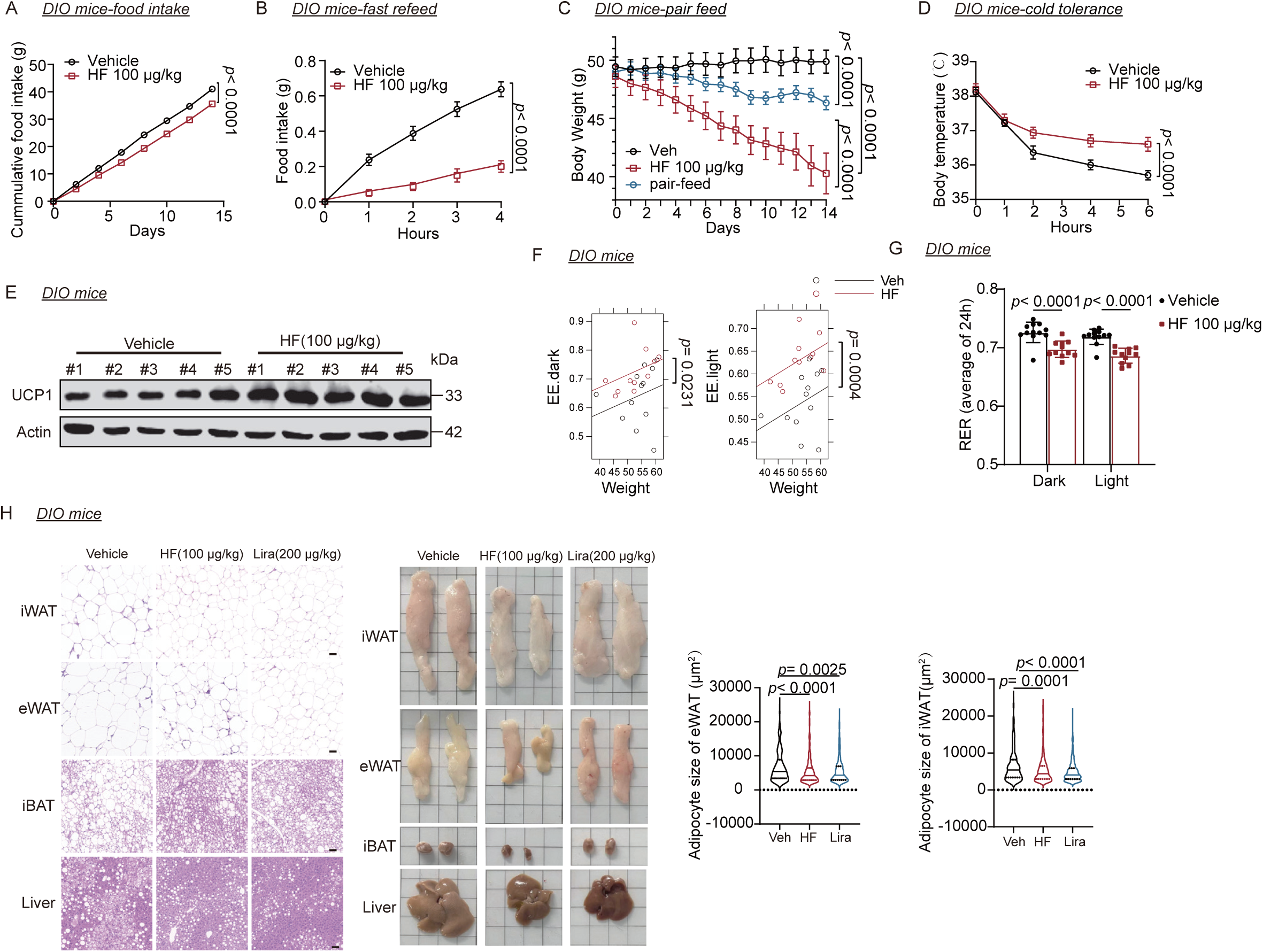
Halofuginone suppresses food intake and increases energy expenditure. **(A)** Cumulative food intake of DIO mice (single cage) treated with vehicle or HF 100 μg/kg (n=8). **(B)** Food intake after fast-refeed DIO mice treated with vehicle or HF 100 μg/kg (n=8). **(C)** Changes in body weight during pair-feeding (n=8). (**D)** Cold tolerance of DIO mice treated with vehicle or HF 100 μg/kg (n=5). (**E)** UCP1 protein abundance in iBAT of vehicle-or HF (100 μg/kg)-treated DIO mice (n=5). (**F)** Energy expenditure (EE) in DIO mice treated with vehicle or HF 100 μg/kg (vehicle group: n = 12, HF group: n=11). (**G)** Average 24 h respiratory exchange ratio (RER) in DIO mice treated with vehicle or HF 100 μg/kg (vehicle group: n = 12, HF group: n=11). (**H)** Representative images of H&E staining of adipose tissues and liver, and the quantification of adipocyte size of eWAT and iWAT. Data are presented as mean ± s.e.m. Data in (A-D) were determined through two-way ANOVA followed by Bonferroni’s multiple-comparison test. Data in (F) were determined through ANCOVA using body mass as a covariate. Bar graphs in (G) analyzed by one-way ANOVA followed by Bonferroni’s multiple-comparison test. Data in (H) were analyzed by non-parametric tests.

HF-treated mice also exhibited reduced weights of inguinal white adipose tissue (iWAT) (-35.6%) and epididymal white adipose tissue (eWAT) (-45.7%), accompanied by consistently smaller adipocyte size and diminished adipose depots (Fig.2H, Extended Data Fig. 3D-E). Compared to the vehicle group, iBAT weight was decreased in HF-treated obese mice, but the iBAT had less lipid accumulation (Fig. 2H, Fig. S3F). Additionally, HF significantly reduced liver weight and ameliorated hepatic steatosis as measured by hepatic TG and TC levels and histological examination, suggesting that HF ameliorated obesity-related fatty liver (Fig.2H, Fig. S3G-I).

### Halofuginone induces GDF15 and FGF21

To unravel the potential molecular mechanisms by which HF suppressed food intake and elevated energy expenditure, we performed bulk RNA sequencing on white adipose tissue and liver of DIO mice (Fig.3A-C). We intersected the differentially expressed genes in white adipose tissue and liver, identifying 55 genes with commonly upregulated expression and 75 genes with commonly downregulated expression (Fig.3D-E). Consistent with previous reports in other tissues ^24^, we observed that HF activated the GCN2/ATF4 signaling pathway in mouse liver (Fig.3F). In earlier reports, ATF4 has been identified as a transcription factor that responds to the ISR and is capable of enhancing the expression of GDF15 and FGF21^25–28^. GDF15 and FGF21, in turn, participate in regulation of energy balance through their respective mechanisms of action. Based on previous research findings and our sequencing results, we focused on two identified hormones, GDF15 and FGF21, hypothesizing that they play key roles in the weight loss process induced by HF. Given that both FGF21 and GDF15 function as secretory proteins, there is ongoing debate about the primary organs that secrete GDF15, with the liver, ileum and kidney believed to be the main sources^25,27,29^. FGF21, on the other hand, is reported to be predominantly secreted by the liver^30^. Hence, we focused on the effects of HF on the GCN2/ATF4 signaling and the expression of GDF15 and FGF21 in the liver and hepatocytes. In DIO mice treated long-term with HF, there was an increase in ATF4 expression (Fig.3G). Concurrently, circulating levels of GDF15 and FGF21 were elevated (Fig.3H). However, HF did not affect circulating levels of the other energy homeostasis-related hormones, including leptin and adiponectin (Fig.3H).

**Fig. 3.**
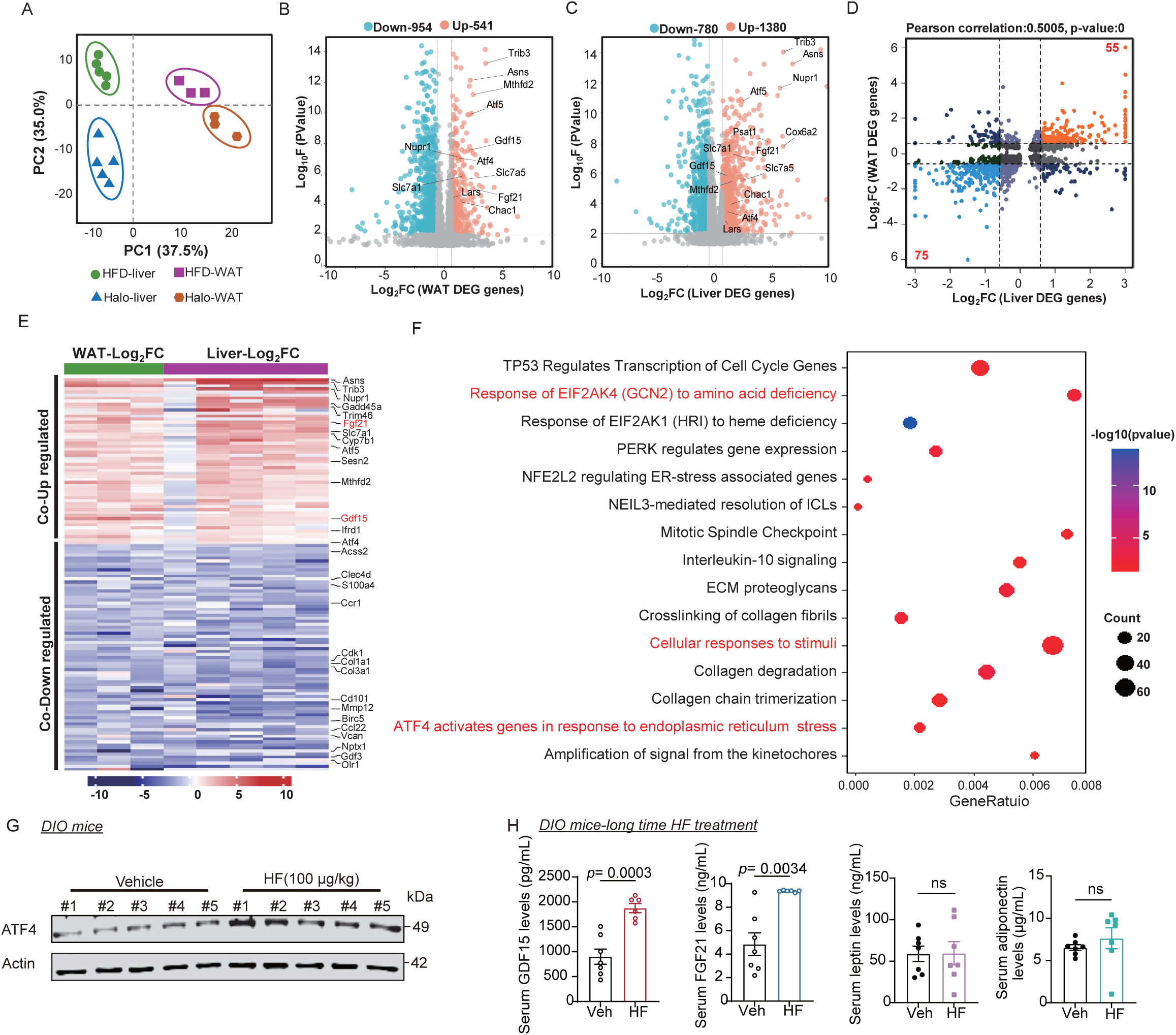
Halofuginone induces GDF15 and FGF21. **(A)** PCA from RNA-seq of vehicle- or HF (100 μg/kg)-treated mouse liver and WAT tissues (WAT: n = 3, Liver: n=5). **(B)** Volcano plot of significantly downregulated (blue) and upregulated (red) genes in WAT from mice as described in panel A (|Log2FC|>1.5, *P* value<0.01). **(C)** Volcano plot of significantly downregulated (blue) and upregulated (red) genes in liver from mice as described in panel A (|Log2FC|>1.5, *P* value<0.01). **(D)** The nine-quadrant plot shows the correlation between differentially expressed proteins in the WAT and liver groups. **(E)** Heatmap analysis of differentially expressed genes in livers and WAT from vehicle- and HF-treated mice. **(F)** Reactome enrichment pathway analysis implicates GCN2/ATF4-associated cellular responses to stress pathway in HF treated group. Significantly over-represented pathways (FDR < 0.05) were grouped and depicted. The size of the circles corresponds to the number of genes in each module. **(G)** Levels of ATF4 protein in liver tissues of DIO mice injected with vehicle or HF (100 μg/kg), n=5. **(H)** Serum GDF15 (vehicle group: n = 7, HF group: n=6), FGF21 (vehicle group: n = 7, HF group: n=6), leptin (n=7) and adiponectin (n=7) protein levels of DIO mice injected with HF (100 μg/kg) for 8 week. Data are meanLJ±LJs.e.m. *P* values for the data of GDF15 and FGF21 were calculated by two-sided unpaired t-tests. *P* values for the data of leptin were calculated by two-sided unpaired t-tests. *P* values for the data of adiponectin were calculated by two-sided unpaired t-tests with Welch’s correction.

### Halofuginone elevates GDF15 and FGF21 expression via the GCN2/ATF4 signaling pathway

To determine whether HF can elevate GDF15 and FGF21 levels before weight loss effects occur, DIO mice were given short-term injections of HF (at intervals of 3 hours, 6 hours, and 12 hours). Our data demonstrate increased circulating levels of GDF15 and FGF21 post acute HF treatment, with GDF15 levels peaking at 3 hours (Fig.4A-B). Moreover, we detected the expression of *Gdf15* and *Fgf21* genes in various tissues (including kidney, liver, ileum, iWAT, eWAT and BAT) of mice following a single intraperitoneal administration of vehicle or HF after 1 h (Fig.4A-B). Compared to the control group, in the liver of the HF group, an upregulation of *Gdf15* (fold change = 11) and *Fgf21* (fold change = 15.6) was observed in the HF group (Fig.4C). Additionally, in Huh-7 cells, HF induced the upregulation of *ATF4*, *GDF15* and *FGF21* gene expression as well as p-GCN2/p-eIF2a/ATF protein expression. However, the upregulation of ATF4, GDF15 and FGF21 as well as GCN2 pathway was blocked by proline supplementation, indicating amino acid response do play an essential role in HF mediated pharmacological effects (Fig.4D-G). Further, we demonstrated through pharmacological inhibition of GCN2 (by applying a pharmacological inhibitor of GCN2, GCN2ib) and genetic inhibition of ATF4 (by applying an siRNA against ATF4), that HF induces the upregulation of *GDF15* and *FGF21* via the GCN2/ATF4 signaling pathway. Thus, ATF4 is identified as the primary transcription factor mediating the effects of HF on elevating GDF15 and FGF21(Fig.4H-I).

**Fig. 4.**
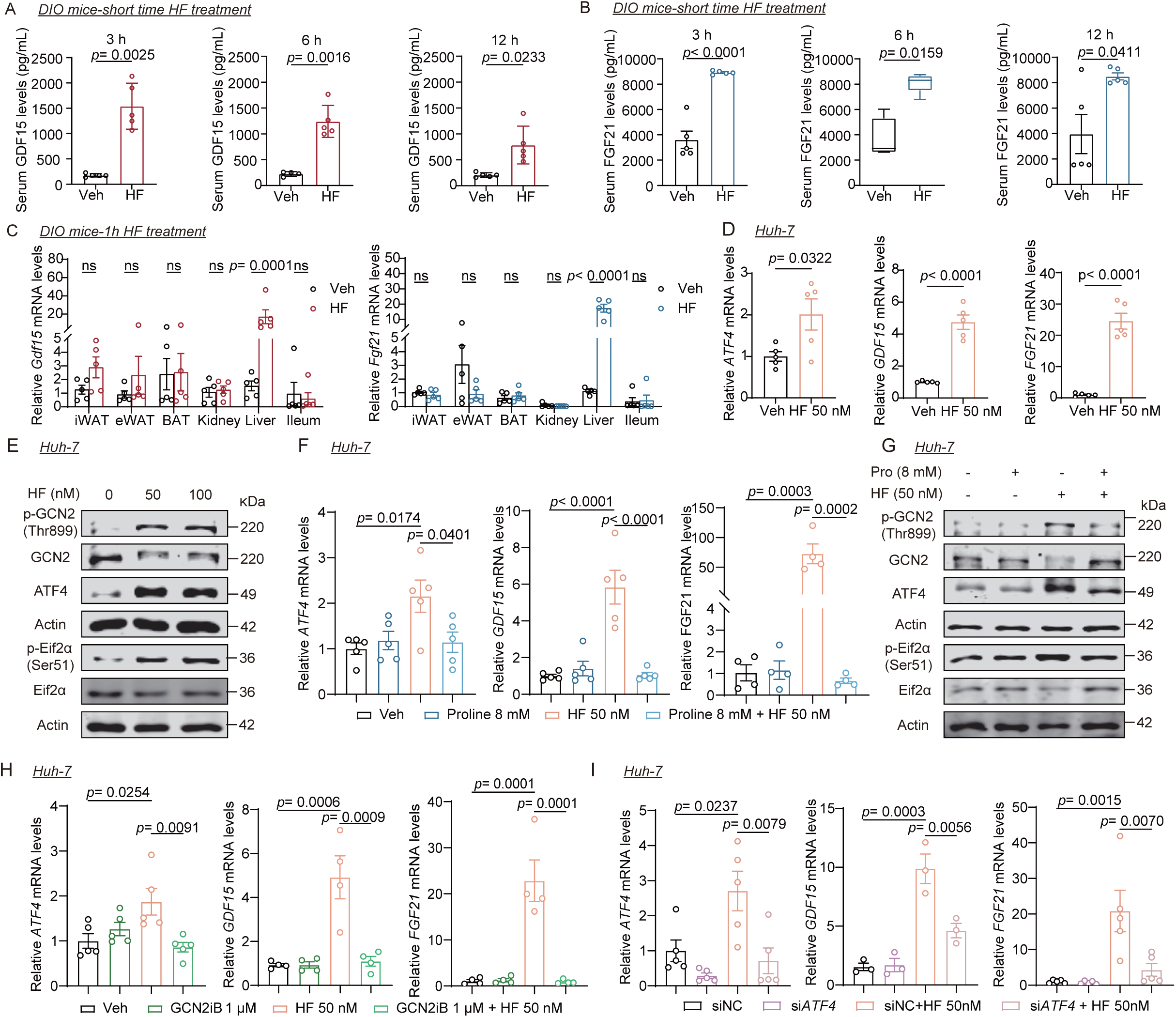
Halofuginone elevates GDF15 and FGF21 expression via the GCN2/ATF4 signaling pathway. **(A)** Serum GDF15 protein levels of DIO mice injected with HF (100 μg/kg) at indicated time (n=5, except for the vehicle group at 6 h (n=4)). **(B)** Serum FGF21 protein levels of DIO mice injected with HF (100 μg/kg) at indicated time (n=5, except for the vehicle group at 6 h (n=4)). **(C)** Levels of *Gdf15* and *Fgf21* mRNAs in indicated organs of DIO mice injected with vehicle or HF (100 μg/kg) at 1 h. **(D)** mRNA levels of *ATF4*, *GDF15* and *FGF21* in human hepatocyte cell line (Huh-7 cells) treated with HF (50 nM) for 24 h (n = 5). **(E)** Protein levels (p-GCN2, GCN2, ATF4, p-eif2α, eif2α) in Huh-7 cells. **(F**) Levels of *ATF4*, *GDF15* and *FGF21* mRNAs in Huh-7 cells treated with HF (50 nM) in the presence or absence of proline (8mM) for 24 h (n = 4-5). **(G**) Protein levels (p-GCN2, GCN2, ATF4, p-eIF2α, eIF2α) in Huh-7 cells. **(H)** Levels of *ATF4*, *GDF15* and *FGF21* mRNAs in Huh-7 cells treated with HF (50 nM) in the presence or absence of GCN2ib (GCN2 inhibitor, 1 μM) for 24 h (n = 4-5). **(I**) Levels of *ATF4*, *GDF15* and *FGF21* mRNAs in control siRNA (siNC) or ATF4 siRNA (siATF4)-treated Huh-7 cells exposed to HF (50 nM) (n =3-5). Data are presented as mean ±LJs.e.m. Data in (A), 3h and 12h in (B) and (D) were calculated using two-sided unpaired t-tests. Data in 6h in (B) were analyzed by non-parametric tests. Bar graphs in (C) analyzed by one-way ANOVA followed by Bonferroni’s multiple-comparison test. Data in (F), (H) and (I) analyzed by one-way ANOVA followed by Bonferroni’s multiple-comparison test.

### GDF15 mediates the food intake suppression and weight-loss effect of HF

To test the involvement of GDF15 in mediating the effects of HF on weight reduction, we utilized *Gdf15* knockout (KO) mice. Circulating levels of GDF15 in KO mice were undetectable, confirming successful knockout of *Gdf15* (Fig.S4A). Ablation of *Gdf15* reversed the terminal weight-loss effect of HF by 27% and fat mass by 55% (Fig.5A-B). Furthermore, HF-induced reductions in iWAT mass in WT mice were reversed in *Gdf15* KO mice, consistent with the observed reduction in body weight (Fig.5C). While HF treatment led to a decrease in plasma levels of triglycerides (TG) and total cholesterol (TC), these favorable effects were not observed in the absence of *Gdf15*, suggesting a pivotal role for *Gdf15* in mediating these impacts (Fig.5D). Additionally, the capacity of HF to reduce food intake was completely abolished in the absence of circulating GDF15 (Fig.5E). Importantly, *Gdf15* KO did not alter the effects of HF on the upregulation of UCP1 and energy expenditure (Fig.5F-G). Additionally, while HF administration improved glucose tolerance and insulin sensitivity in WT mice, these metabolic benefits of HF were also evident in *Gdf15* KO mice, suggesting, importantly, that *Gdf15* is not involved in mediating the effects of HF on glucose intolerance and insulin sensitivity (Fig. S4B-C). Collectively, these findings highlight the essential role of *Gdf15* in HF-mediated effects on obesity in mice, while still indicating the existence of other mechanisms contributing to the metabolic benefits of HF.

**Fig. 5.**
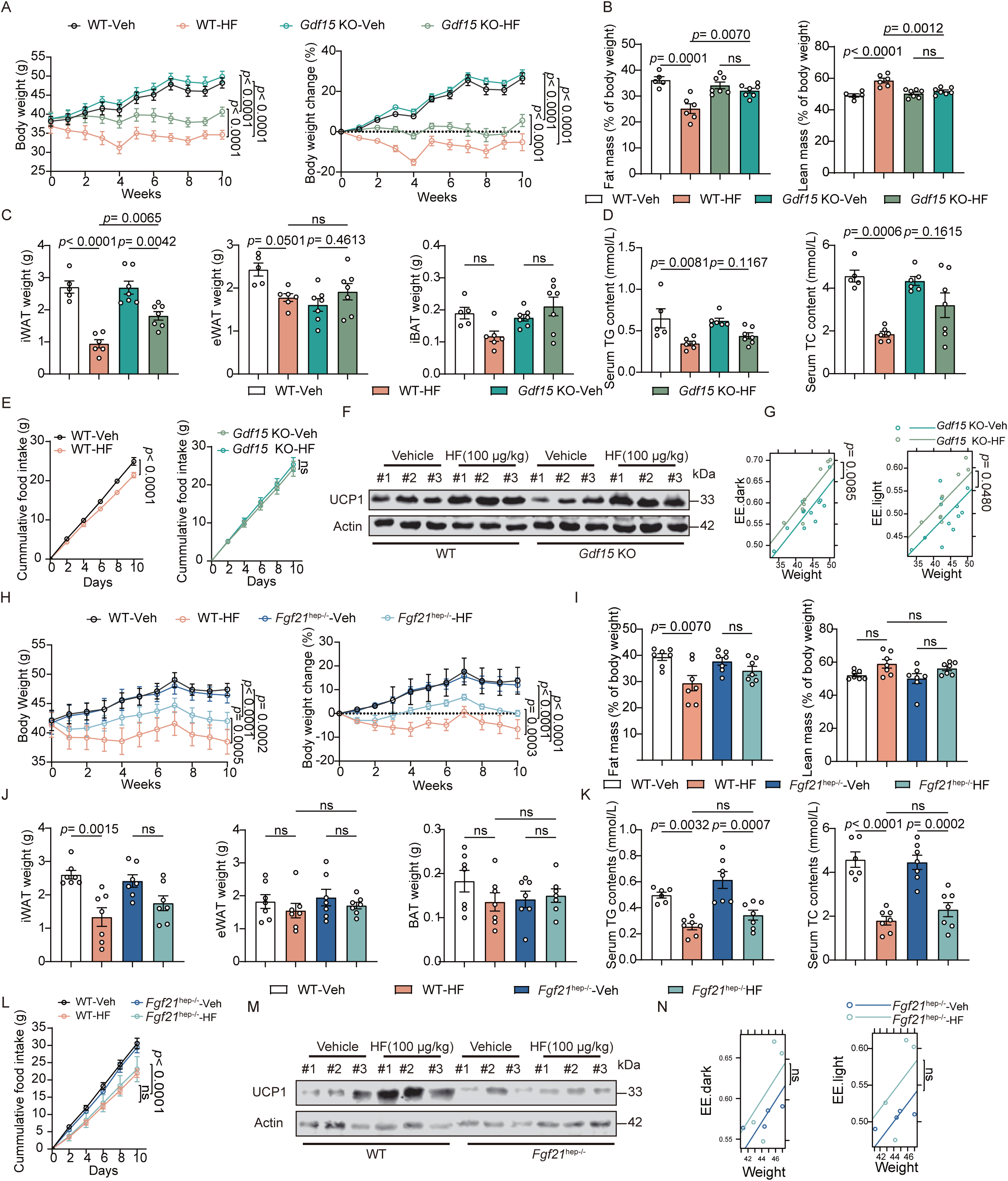
GDF15 and FGF21 mediates weight-loss effect of Halofuginone by inhibiting food intake and increasing energy expenditure, respectively. **(A-D)** 8-week-old male wild-type (WT) and *Gdf15*^-/-^ mice were fed with HFD for 8 weeks. WT and *Gdf15*^-/-^ mice were then randomly assigned to vehicle or HF (100 μg/kg) group and injected every two days for 8 weeks. The mice were subjected to the following measurements: **(A)** Body weight and % body weight change (WT-Veh: n = 5, WT-HF: n=6, *Gdf15* KO-Veh: n=7, *Gdf15* KO-HF: n=7). **(B)** Fat mass and lean mass (% body weight). **(C)** Weights of adipose tissues. **(D)** Serum TC and TG levels. **(E)** Cumulative food intake of single cage WT or *Gdf15* KO mice treated with vehicle or HF 100 μg/kg (vehicle group: n = 10, HF group: n=10). **(H-K)** 8-week-old male *Fgf21*^flox/flox^ mice (WT) and *Fgf21*^hep–/–^ mice were fed with HFD for 12 weeks, and then randomly divided into vehicle or HF (100 μg/kg) group and injected every two days for 10 weeks. **(H)** Body weight and % body weight change (n=7). **(I)** Fat mass and lean mass (% body weight) (n=7). **(J)** Weights of adipose tissues (n=7). **(K)** Serum TC and TG levels (n=7). **(L)** Cumulative food intake of single cage WT or *Fgf21*^hep–/–^ mice treated with vehicle or HF 100 μg/kg (n=4). **(M)** UCP1 protein abundance in BAT of HF (100 μg/kg)-treated WT and *Fgf21*^hep*-/-*^ mice (n=3). **(N)** EE in *Fgf21*^hep*-/-*^ mice treated with vehicle or HF 100 μg/kg (vehicle group: n = 4, HF group: n= 4). Data are presented as mean ±LJs.e.m. Data in (A), (E), (H), and (L)were determined through two-way ANOVA followed by Bonferroni’s multiple-comparison test. Data in (G) and (N) were determined through ANCOVA using body mass as a covariate. Bar graphs in (B-D) and (I-K) analyzed by one-way ANOVA followed by Bonferroni’s multiple-comparison test.

### FGF21 mediates the increased energy expenditure and weight-loss effect of HF

We conducted further investigations to ascertain whether or not hepatic FGF21 mediates the additional effects of HF on weight loss. We established hepatocyte-specific *Fgf21* knockout mice (*Fgf21*^hep-/-^ mice) through the crossbreeding of *Fgf21* ^flox/flox^ mice (WT mice) with Alb-cre mice, followed by administration of HF to both WT and *Fgf21*^hep-/-^ mice. Circulating FGF21 levels were reduced to below 110 pg/mL (90% decrease) in hepatocyte-specific FGF21^hep-/-^ mice, thereby confirming successful hepatocyte *Fgf21* deletion (Fig.S5A). Ablation of hepatic *Fgf21* reversed the terminal weight-loss effect of HF by 44% and that of fat mass by 53% (Fig.5H-I). Consistent with findings observed in *Gdf15* KO mice, HF administration reduced iWAT mass in WT mice, however, the effect of HF on iWAT loss was not observed in *Fgf21*^hep-/-^ mice (Fig.5J). Consistent with the reduction in body weight, HF treatment resulted in reduced plasma TG and TC levels. However, *Fgf21* hepatic deletion showed minimal impact on the lipid-lowering effect of HF (Fig.5K). Notably, hepatic ablation of FGF21 did not reverse the ability of HF to reduce food intake (Fig.5L), but significantly inhibited the effects of HF on upregulation of UCP1 and HF-elicited increase in energy expenditure (Fig.5M-N). HF induced ameliorative effects on glucose intolerance and insulin resistance was mildly affected by hepatocyte-specific deletion of *Fgf21* (Fig.S5B-C). Collectively, these findings demonstrate the obligatory role of hepatocyte-derived *Fgf21* in the HF-mediated impact on weight loss.

## Discussion

In the present study, we identified HF, a derivative of the anti-malarial quinazolinone natural product alkaloid, febrifugine, as an inducer of metabokines GDF15 and FGF21, and a regulatory agent for food intake and energy expenditure and hence body weight. This pharmacological, metabolic and mechanistic data supports the potential of HF for treating obesity and its comorbidities. Previous studies have shown that dietary protein restriction or reduction of specific amino acids, such as methionine and branched-chain amino acids (BCAAs), promote metabolic health through pathways including GCN2/ATF4, mTOR, and AMPK^12,13^. The AAS response, triggered by endogenous amino acid depletion, activates metabolic regulation mechanisms that remain understudied. In this regard, inhibition of EPRS by HF leads to the accumulation of uncharged tRNAs, initiating autophosphorylation of the amino acid sensor GCN2, resulting in the phosphorylation of eIF2α and increased expression of ATF4. The involvement of the GCN2 pathway in various metabolic processes has been already documented ^31,32^.

Obesity and related metabolic abnormalities are commonly attributed to energy imbalance due to excess food (calorie) intake and/or insufficient physical activity ^1,2^. The therapeutic potential of GDF15 and FGF21 in obesity and metabolic liver diseases has been explored in preclinical studies and clinical trials ^33–36^. For example, the weight loss efficacy of metformin relies on GDF15^29,37,38^. A long-acting GDF15 analog showed promising results in obese Cynomolgus monkeys, leading to 16 ± 5% weight loss^39^. In mice, GDF15 stimulates the sympathetic nervous system, leading to increased calcium release and energy expenditure in muscle^40^. This dual action on appetite suppression and metabolic rate makes GDF15 a compelling target for weight loss therapies. More recently, artesunate has been reported to treat obesity in mice and non-human primates through GDF15/GFRAL signalling axis^41^. Beyond GDF15, FGF21 analogues have also shown robust effects on lipid metabolism and liver function, rendering them as promising candidates for treating metabolic disorders, including MASH. The Phase 2 clinical trial results for pegozafermin, an FGF21 analog, indicated that, after 24 weeks, the highest dose group demonstrated a one-stage or better improvement in liver fibrosis^42^. In this study, we observed that HF rapidly increased serum levels of GDF15 and FGF21, mediated by the canonical GCN2/ATF4 pathway. *In vitro*, the supplementation of proline in hepatocytes negated the HF-induced increase in GDF15 and FGF21. Further, both GCN2 inhibition or ISR inhibition suppressed the elevation of GDF15 and FGF21 induced by HF. This suggests that HF acts through inhibition of EPRS and simulation of the AAS pathway to activate ISR, leading to elevated circulating levels of both GDF15 and FGF21. *In vivo* data obtained with MAZ1310 treatment of obese mice provided additional support for the involvement of EPRS inhibition in HF-induced weight loss effects. Previous work shows that GDF15 reduces appetite, enhances energy expenditure in muscle, and thereby reduces body weight ^29,40,43^, while FGF21 administration enhances energy expenditure, boosting insulin sensitivity and resulting in weight loss, partially independent of effects on food intake ^44,45^. However, clinical trials using protein-based therapies face challenges in improving metabolic disorders due to protein stability, administration methods, and patient adherence to treatment regimes. As a potential therapeutic agent, GDF15 encounters substantial obstacles in terms of druggability. For example, GDF15 has a very short half-life, approximately 3 hours in both mice and Cynomolgus monkeys^46^. Moreover, GDF15 exhibits a strong propensity to aggregate, leading to low stability and expression titers. In contrast, HF has significant advantage and effectively acts as an endogenous mimetic by stimulating endogenous secretion of GDF15 and FGF21 and maintaining elevated circulating levels of both metabolic hormones for extended periods. This is a novel approach for the therapy of obesity-associated metabolic disorders based on the simultaneous action of these two proteins stimulated by a single drug. In addition to HF, other intervention strategies, such as iD1^47^ (a pharmacological inhibitor of CNOT6L deadenylase), and dietary intervention (ketogenic diet)^48^, have been reported to activate GDF15 and FGF21, leading to weight loss and amelioration of metabolic disorders. These findings are in harmony with our findings. ID1 enhances the levels of GDF15 and FGF21 through inhibiting the action of CNOT6L^47^, while the ketogenic diet primarily exerts its beneficial effects via hepatic PPARγ^48^. This convergence of different mechanisms underscores the pivotal roles of GDF15 and FGF21 in maintaining energy homeostasis and metabolic health. Despite targeting different pathways, these interventions achieve similar outcomes, highlighting the versatility and importance of GDF15 and FGF21 in addressing metabolic diseases.

We demonstrate here the weight loss efficacy of HF in both diet-induced and genetically modified obese mouse models. Since humans live at thermoneutrality, drug testing using mouse models are suggested to include experiments performed under near thermoneutrality (TN) conditions^49^. Our data suggested that HF also leads to weight loss in DIO mice under TN conditions, further supporting the clinical translatability of HF. Notably, we also observed that HF-induced weight loss benefits extend to oral route, further extending its potential utility and acceptability in clinical settings. Also, its effects were consistent in both male and female mice, indicating a stable and reliable weight loss outcome irrespective of sex. Moreover, in *ob/ob* mice, HF treatment exhibited appreciable weight loss effects in this genetic-relevant animal model. These evidences collectively highlight the robust weight-loss capabilities of HF, and its efficacy across different routes of administration, sexes, and animal models under different housing temperatures, underscoring its potential as a versatile and effective agent for managing obesity.

HF shows potential for translation into an anti-obesity treatment. Previous studies have reported HF’s anti-malarial, anti-inflammatory, anti-fibrotic, anti-cancer, and anti-viral effects, supporting its clinical translational value due to its polypharmacological actions^15–18^. Clinical trials of halofuginone have centered on the evaluation of its optimal dosage and therapeutic windows for HIV-related kaposi’s sarcoma and advanced solid tumors (www.clinicaltrials.gov). In phase I and II clinical trials, the recommended oral dose of HF for long-term administration is 0.5 mg/day^19,20^. In our experiments, the dose of HF recommended for anti-obesity purposes (100 µg/kg) is approximately half of the lowest dose tested in phase II clinical trials, and even lower doses of HF (25 µg/kg or 50 µg/kg) have been effective in controlling body weight in mice. In non-obese animals fed a standard diet, HF administration had no significant adverse effects in major tissues, another assurance of its safety.

Although HF improved the metabolic traits of obese mice, caution must be exercised as pharmacologically modulating AAS pathways may have broad impacts on cellular functions beyond metabolism. In addition, given the broad-spectrum effects of HF, the primary tissue source of circulating GDF15 during HF treatment were not thoroughly explored. Our data suggest the liver may be a dominant source of circulating GDF15, which warrants further verification in liver-specific *Gdf15* deficient mice. Thus, mice deficient of both hepatic *Gdf15* and *Fgf21* are needed to clarify the full-spectrum of anti-obese effects of HF.

In summary, we report HF as a previously unexplored therapeutic agent to ameliorate diet-induced metabolic disorders through orchestrating food intake and energy expenditure via dual elevation of GDF15 and FGF21. This discovery opens up avenues for controlling obesity by dual targeting of GDF15 and FGF21. Future efforts will be required to translate the therapeutic benefits of HF to obese subjects, as well as define the long-term safety profile of activating AAS response pharmacologically.

## Acknowledgments

The authors are grateful to Profs. Jian Liu and Xian Zhang (Hefei University of Technology, Hefei, China), Prof. Cheng Zhan (University of Science and Technology of China, Hefei, China) and Prof. Ze Zheng (Medical College of Wisconsin, Milwaukee, USA) for insightful discussions. We thank members of the Weng laboratory for valuable discussions and feedback. S.X. is Senior Humboldt Research Fellow of Alexander von Humboldt Foundation, Germany.

## Author contributions

Conceptualization: J.W., S.X. and J.R.S. Investigation: Z.L., T.T., W.Z., M.L., M.X., M.C., M.S., F.Z., W.P., Z.Z., Z.W., S.L. and Y.Y. Funding acquisition: J.W., S.X. Writing – original draft: Z.L., T.T., S.X. Writing – review & editing: P.J.L., D.K., D.W., J.W., M.D.L., S.Z., S.O. and J.R.S.

## Data and materials availability

All data generated and used in this study are either included in this article (and its Supplementary Information) or are available from the corresponding author on reasonable request. Data will be made available on request. RAW data of RNA-seq was deposited in Gene Expression Omnibus with accession number: GSE273929.

## Supplementary Materials

Materials and Methods

Figs. S1 to S5

