## supplemental materials for "The clinical antiprotozoal drug halofuginone promotes weight loss by elevating GDF15 and FGF21"

**The PDF file includes:**

Materials and Methods
Figs. S1 to S5

### Materials and Methods

#### Animal studies

All experimental procedures were conducted in accordance with the "Guide for the Care and Use of Laboratory Animals". Approval for the animal experiments was granted by the Institutional Animal Care and Use Committee of University of Science and Technology of China (Approval number: USTCACUC27120124062). Eight-week-old C57BL/6J mice (Strain NO. N000013) and *ob/ob* mice (Strain NO. T001461) were obtained from GemPharmatech (Nanjing, China). *Gdf15*$^{-/-}$ (Strain NO. T011862)[50,51] and *Alb-cre* (Strain NO. T003814) mice were purchased from GemPharmatech (Nanjing, China). *Fgf21*$^{flox/flox}$ (NM-CKO-00136) mice were purchased from Shanghai Model Organisms Center (Shanghai, China). Mice were housed in a specific-pathogen-free (SPF) environment with 12 h of light and 12 h dark cycle and were maintained at 23 ± 2°C with free access to food and water. Animals were randomly divided into different groups before subject to high fat-det diet (HFD, #D12492, Research Diets, NJ, USA).

#### Drug treatment

For HF treatment, mice received different doses of HF (Targetmol, T3524, China) or vehicle (0.1% dimethyl sulfoxide mixed with 0.9% physiological saline) intraperitoneally (*i.p.*) according to the different experimental requirements until the end of the experiment.

#### Animal experiments

**Mouse experiment 1: effect of HF on DIO mice**

Mice (8-week-old male C57BL/6J) were fed a HFD for 16 weeks, and then randomly assigned to four experimental groups: (1) vehicle group, (2) 25 μg/kg HF treatment group, (3) 50 μg/kg HF treatment group or (4) 100 μg/kg HF treatment group, with five mice per group, (5) 200 μg/kg liraglutide (a GLP1R agonist, reference drug) treatment group with four mice in group. During HFD challenge, mice were treated with intraperitoneal injections of vehicle and HF every two days. The body weights were monitored weekly. After an 8-week injection period, mice were euthanized after starvation overnight.

**Mouse experiment 2: HF withdrawal experiment**

Mice (8-week-old male C57BL/6J) on a HFD for 12 weeks were randomly divided into two groups: (1) vehicle group (n=10) or (2) 100 μg/kg HF treatment group (n=20). The mice were injected with vehicle and HF every two days for 6 weeks, then the HF group was randomly divided into two groups:(1) continuous administration group (n=10), (2) withdrawal group (n=10). The body weights were monitored weekly.

**Mouse experiment 3: effect of HF on *ob/ob* mice**

Male *ob/ob* mice (8-week-old) were raised under a standard chow diet and randomly divided into 2 groups: (1) vehicle (n=7) or (2) 100 μg/kg HF treatment group (n=8). Mice were treated with intraperitoneal injections of vehicle and HF every two days. Animals were weighted and weights recorded every week. At the end of 8-week testing, mice were euthanized after starvation overnight.

**Mouse experiment 4: effect of HF on lean mice**

To monitor the chronic safety of long-term treatment with HF, male 8-week-old C57BL/6J mice were injected intraperitoneally with vehicle or 100 μg/kg HF every two days for 28 weeks. Body weights were monitored every weekly. At the end of experiment, mice were euthanized by anesthesia after starvation overnight.

**Mouse experiment 5: effect of oral HF on DIO mice**

Mice (8-week-old male C57BL/6J) were randomly divided into 5 groups: (1) Normal diet control, (2) HFD control, (3) HFD+100 μg/kg HF, (4) HFD+200 μg/kg HF, (5) HFD+20 mg/kg orlistat (an orally bioavailable anti-obese reference drug). After 12 weeks of normal diet or HFD feeding, the mice were treated with vehicle or HF every two days by oral gavage.

**Mouse experiment 6: effect of HF on female mice**

Mice (8-week-old female C57BL/6J) were fed HFD for 20 weeks, and then randomly assigned for two groups: (1) vehicle group (n=5), (2) 100 μg/kg HF treatment group (n=5). The body weights were monitored weekly.

**Mouse experiment 7: pair feeding**

Mice (8-week-old male C57BL/6J) with HFD for 12 weeks were randomly divided into three groups: (1) vehicle group (n=8), (2) 100 μg/kg HF treatment group (n=8), (3) pair-fed vehicle group (n=8). Pair-fed vehicle-treated mice received the same amount of food as ingested by the corresponding HF-treated groups the day before. Body weight and food intake were recorded daily.

**Mouse experiment 8: short-term HF treatment**

Mice (8-week-old male C57BL/6J), fed HFD for 12 weeks, were euthanized at 1 h, 3 h, 6 h and 12 h post a single bolus treatment with vehicle or HF.

**Mouse experiment 9: effect of HF in *Gdf15*$^{-/-}$ mice**

8-week-old male C57BL/6J mice (wild-type) and *Gdf15*$^{-/-}$ mice were fed with HFD for 8 weeks. Wild-type and *Gdf15*$^{-/-}$ mice were then randomly assigned to vehicle or HF (100 μg/kg) group and injected every two days for 8 weeks. Mice were monitored weekly for body weight and food intake, and then were sacrificed by anesthesia after starvation overnight after 12 weeks of HF treatment.

**Animal experiment 10: effect of HF in *Fgf21*$^{hep-/-}$ mice**

*Fgf21*$^{hep-/-}$ mice were obtained by were crossing *Fgf21*$^{flox/flox}$ mice with *Alb-cre* mice. 8-week-old male *Fgf21*$^{flox/flox}$ mice (wild-type) and *Fgf21*$^{hep-/-}$ mice were fed with HFD for 12 weeks, and then randomly divided into vehicle or HF (100 μg/kg) group. The mice were injected intraperitoneally every two days for 11 weeks and were sacrificed at the end of experiment.

**Fasting-refeeding protocol**

The fasting-refeeding studies were performed in 12-week HFD mice. Before the experiment began, mice were acclimated to single cage for 48 h, and then mice were subjected to 16 h fasting. Food was returned to cages after 1 h of HF treatment. The food intakes ware measured at 1, 2, 3, 4 h after refeeding.

**Pharmacokinetics and tissue distribution of HF**

For pharmacokinetic studies, after a single intravenous (168 μg/kg) or oral administration (840 μg/kg) of HF in ICR (CD-1) mice (n = 3 per group), plasma samples were collected at indicated time points (0.25, 0.5, 1, 2, 4, 8 and 24 h) to measure HF concentrations by high-performance liquid chromatography (HPLC). For tissue distribution studies, after a single intravenous (168 μg/kg) or oral administration (840 μg/kg) of HF in ICR (CD-1) mice, the mice were sacrificed and their plasma and tissues were collected at indicated time points (0.25, 2, 4, and 8h) to measure HF concentrations.

**Cold exposure test**

For cold exposure, mice were exposed to 6°C to test their cold tolerance. The core body temperatures were recorded by a rectal thermometer (Physitemp, Clifton, NJ) at 0 h, 1 h, 2 h, 4 h and 6 h. Mice had free access to water, but not to food.

**Glucose tolerance test and Insulin tolerance test**
For glucose tolerance test, mice were fasted overnight (17:00pm-9:00am) and received intraperitoneal injection of D (+)-glucose (Diamond, A100188, Shanghai, China) solution in saline (2 g/kg body weight). Blood glucose levels were measured from the tail vein by glucometer (Vivachek, VGM83, Hangzhou, China) at 0, 15, 30, 60, 90 and 120 min after injection. For insulin tolerance test, mice were intraperitoneally injected with 0.75U/kg of insulin (Novorapid, Insulin Aspart Injection, Bagsværd, Denmark) after 4h fast (9:00am-13:00pm). Blood glucose levels were measured at 0, 15, 45, 30, 60, 90 and 120 min.

**Blood tests**
Blood was collected and placed in anticoagulant tubes (50 µL) and EP tubes (about 600 µL) respectively. Fresh blood in the anticoagulant tubes was immediately analyzed for blood cell content using an automated blood analyzer (Mindray, BC-30 Vet, Shenzhen, China). The blood in the EP tube were left at room temperature for two hours, centrifuged at 2000×g for 15 min, and the serum was collected and stored in the refrigerator at -80°C.The serum parameters include ALT, AST, TG, TC, HDL, LDL, UREA, CREA and CK were measured by using assay kits (Rayto, Shenzhen, China). Levels of mouse serum GDF15 and medium GDF15 released from cultured hepatocytes were measured by Mouse/Rat GDF-15 Quantikine ELISA Kit (R&D Systems, MGD150, MN, USA). Levels of mouse serum FGF21 and medium FGF21 released from cultured hepatocytes were assessed by Mouse/Rat FGF21 Quantikine ELISA Kit (R&D Systems, MF2100, Minnesota, USA). Levels of insulin were detected by Mouse Insulin ELISA Kit (Mercodia, 10-1247-01, Uppsala, Sweden).

**Body composition measurements**
Total lean mass and fat mass of mice after treatment of either vehicle or HF were assessed using Minispec (Bruker, Massachusetts, USA).

***In vitro* translation experiment with rabbit reticulocytes.**
The Flexi® Rabbit Reticulocyte Lysate System kit from Promega (L4540, WI, USA) was used for the experiments per user guide. In summary, all required components for in vitro translation were assembled in an EP tube, with a final concentration of 200 nM HF, 200 nM MAZ1310, or 8 mM proline. The reaction mixture was incubated at 30°C for 90 minutes, followed by the measurement of luciferase activity.

**Liver TG and TC analysis**
The TG and TC levels of livers were measured by using quantification kits (Wako, 632-50991 and 635-50981, Osaka, Japan.). To perform the quantification, equal weights of liver tissues were first lysed in isopropanol to ensure thorough cell disruption. The homogenate was then centrifuged at 10,000 × g for 10 minutes at 4°C to remove cellular debris. The supernatant was collected and used for the assays.

**Metabolic chamber**

Food intake, oxygen consumption and energy expenditure were determined with CLAMS (Columbus, Ohio, USA). Mice were individually housed and acclimated to the metabolic chambers for three days. Before formal testing, mice were injected intraperitoneally with saline every day for three days. Then vehicle or HF (100 μg/kg) were administered to mice every two days before dark cycle. Hourly data were statistically analyzed using CalR (https://www.CalRapp.org/)[52], and ANCOVA was performed with R Studio.

**Cell culture**
Huh-7 cells (Cellbank, SCSP-526, Shanghai, China) ere cultured in DMEM (KeyGEN biotech, KGL1206-500, Nanjing, China) supplemented with 10% FBS (Sigma-Aldrich, F7524, Missouri, USA). Cells in 12-well plates were used for drug treatment and siRNA knockdown, with a treatment duration of 24 hours. In transfection experiments, cells were treated with drugs for an additional 24 hours following 24 hours of siRNA transfection. At the end of the experiment, cellular RNA and proteins were extracted.

**Histological analysis**
Freshly harvested tissues were promptly fixed in 4% paraformaldehyde overnight to preserve cellular architecture and protein integrity. Following fixation, the tissues were embedded in paraffin wax. The paraffin-embedded samples were then sectioned at a thickness of 5-10 micrometers using a microtome. For histological examination, the tissue sections underwent Hematoxylin and Eosin (H&E) staining.

**Western blotting**
Tissue samples were disrupted utilizing RIPA buffer (Sangon, C500005-0100, Shanghai, China) supplemented with 1× Protease inhibitor (Yeasen, 20124ES03, Shanghai, China) and phosphatase inhibitor (Yeasen, 20109ES05, Shanghai, China) on ice, followed by centrifugation at 12,000×g for 10 minutes. The resulting supernatants were harvested, and the total protein content was determined employing a BCA Protein Assay Kit. Subsequently, a 2 mg/ml protein extract was mixed with loading buffer and subjected to denaturation through boiling at 95°C for 10 minutes. Equal quantities of protein were separated via electrophoresis on either 10% or 12% SDS-page gels and then transferred onto a NC membrane (Pall, PAL-66485, NY, USA). Subsequent incubations with corresponding primary and secondary antibodies were conducted, and imaging was accomplished utilizing the Li-COR two-color fluorescence imaging system. Primary antibodies were as follows: ATF4(Cell signaling technology, 11815, Massachusetts, USA), GCN2(Cell signaling technology, 3302, Massachusetts, USA), p-GCN2(Abcam, ab75836, Cambridge, UK), p-eif2a (Abcam, ab32157, Cambridge, UK), eif2a (Abcam, ab5369, Cambridge, UK), Beta Actin (Proteintech, 66009-1-Ig, Wuhan, China).

**Quantitative PCR**
Total RNA was isolated from tissue samples using the NucleoZol (Gene company, 740404.200, Nanjing, China) according to the manufacturer's protocol. For Huh-7 cells, RNA-Quick Purification Kit (ESScience, RN001, Shanghai, China) were used according to the manufacturer's protocol. The quantity and purity of RNA were determined using a NanoDrop (Thermo Scientific, Massachusetts, USA). Complementary DNA (cDNA) was synthesized from 1 μg of total RNA using PrimeScript™ RT reagent Kit (Takara, RR037A, Kusatsu, Japan) following the manufacturer’s instructions. Quantitative PCR (qPCR) was performed using the

TB Green® Premix Ex Taq™ II (Takara, RR820Q, Kusatsu, Japan) on LightCycler® 96 Instrument (Roche, Basel, Switzerland). Specific primers for target genes (*Atf4*, *Gdf15*, and *Fgf21*) and the reference gene (*Gapdh*) were designed using Primer-BLAST (NCBI).

**Statistical analyses**

Data was expressed as means ± standard error (mean ± s.e.m.) and the statistical significance was set at $p < 0.05$. The statistical analyses were performed with GraphPad Prism10.0 (GraphPad Software, California, USA) and tested by either Student's t-test, one-way or two-way ANOVA as indicated in the figure legends. The sample size and number of replicates for each experiment are described in the figure legends. For "reversed %", we use the following calculation method: the change in value in the WT group after drug administration compared to the vehicle group is subtracted by the change in value in the KO/hep$^{-/-}$ group after drug administration compared to the vehicle group, and then divided by the change in value in the WT group after drug administration compared to the vehicle group.

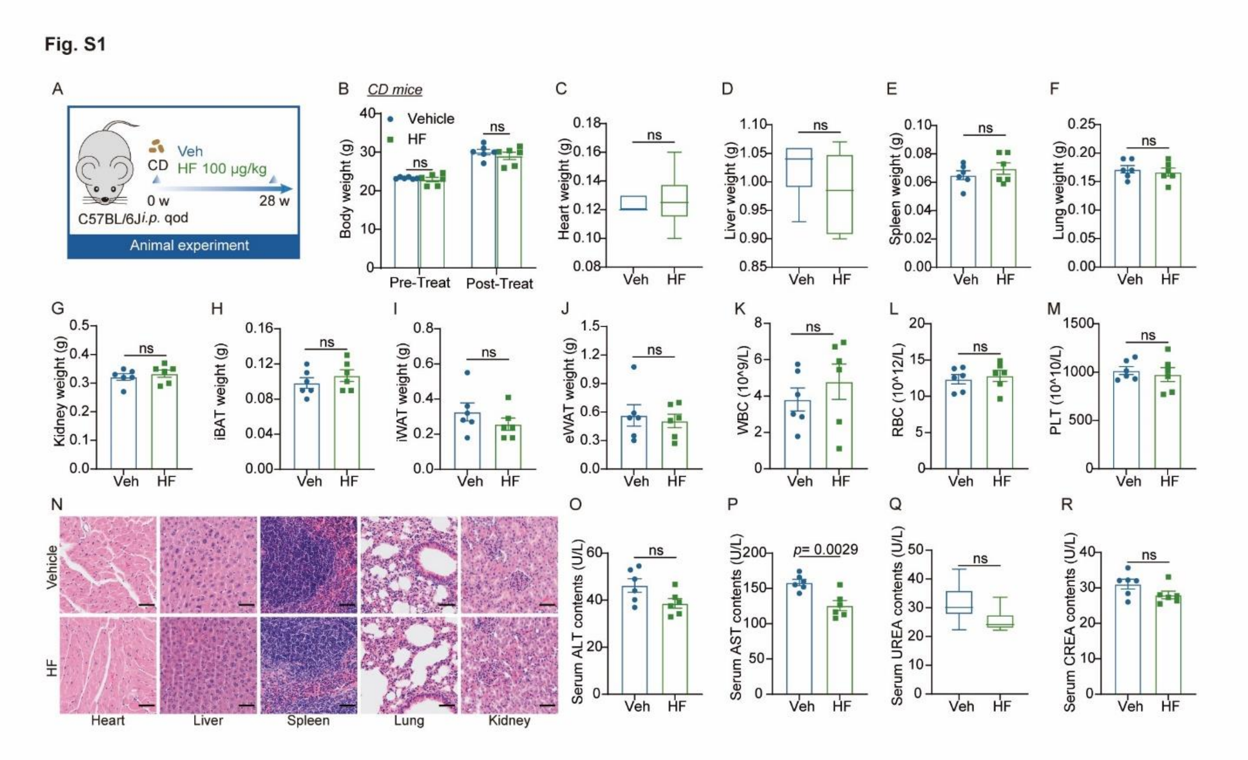


**Fig. S1. Long-term administration of halofuginone is safe in mice.**

**(A-R)** 8-week-old male C57BL/6J mice fed chow diet were injected intraperitoneally with vehicle or 100 μg/kg HF every two days for 28 weeks. **(A)** Schematic diagram illustrating indicated experiment. The mice were subjected to the following measurements: (B) Body weight (n = 6). **(C-J)** Weights of indicated tissues (n=6). **(K-M)** Plasma blood cell components (n=6). (K) WBC: white blood cell, (L) RBC: red blood cell, (M) PLT: platelet. **(N)** Representative H&E staining of dissected tissues. Scale bar, 100 μm. **(O-R)** Serum biochemistry parameters (n=6). Data are presented as mean ± s.e.m. Data in (B) were determined through two-way ANOVA followed by Bonferroni's multiple-comparison test. Data in (C), (D), and (Q) were analyzed by non-parametric tests. Bar graphs in (D-P) and (R) were calculated using two-sided unpaired t-tests.

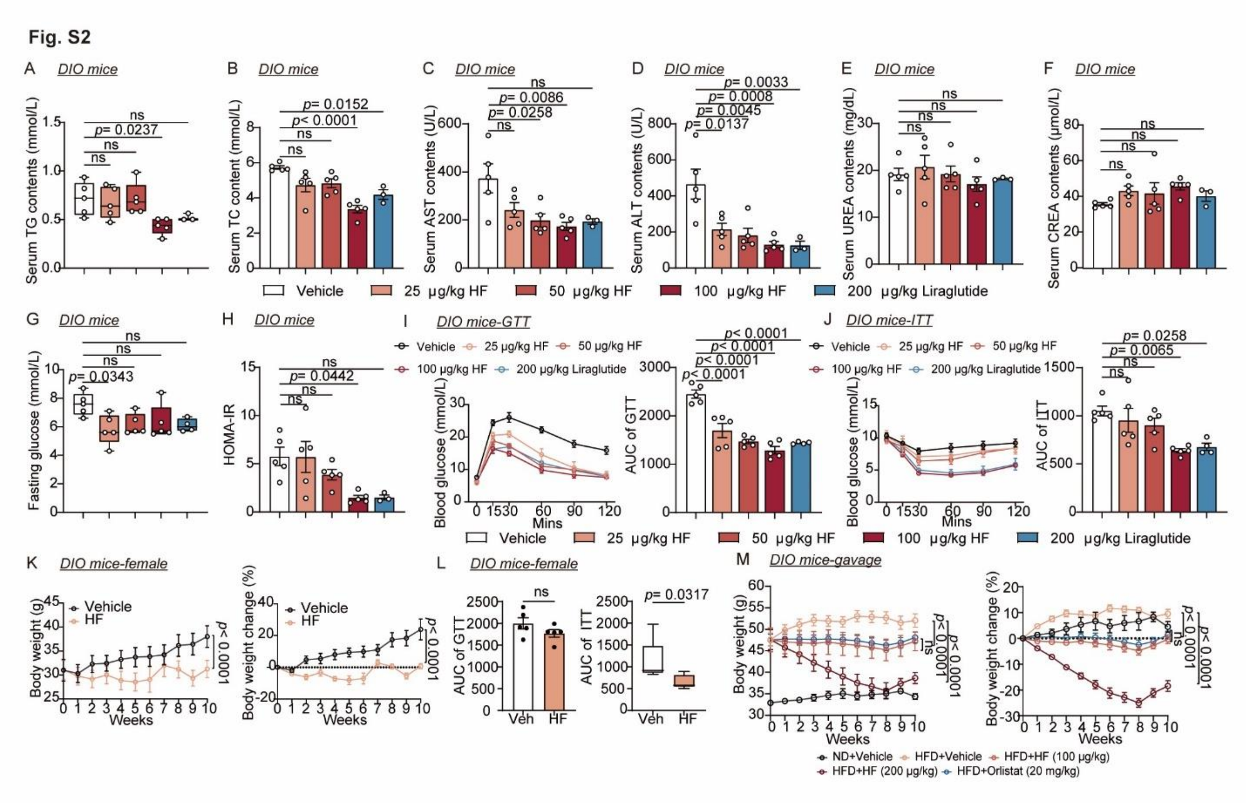


**Fig. S2. Halofuginone ameliorates metabolic dysfunction in mice.**

**(A-J)** 8-week-old male C57BL/6J mice were fed a HFD for 16 weeks, and then randomly assigned to four experimental groups: (1) vehicle group, (2) 25 μg/kg HF treatment group, (3) 50 μg/kg HF treatment group or (4) 100 μg/kg HF treatment group, (5) 200 μg/kg Liraglutide treatment group. During HFD challenge, mice were treated with intraperitoneal injections of vehicle and drugs every two days. The mice were subjected to the following measurements: **(A-F)** Serum parameters (vehicle or HF group: n = 5, liraglutide group: n = 3). **(G-H)** Fasting blood glucose and HOMA-IR (vehicle or HF group: n = 5, liraglutide group: n = 4). **(I-J)** Glucose tolerance test (GTT, I) and insulin tolerance test (ITT, J) (n=5). **(K-L)** 8-week-old female C57BL/6J mice fed high fat diet were injected intraperitoneally with vehicle or 100 μg/kg HF every two days for 10 weeks. The mice were subjected to the following measurements: **(K)** Body weight and % body weight change (n = 5). **(L)** GTT and ITT (n = 5). **(M)** Body weight and % body weight change of male DIO mice treated with vehicle, HF or orlistat by oral gavage (n=5). Data are presented as mean ± s.e.m. Data in (A), (G) and AUC of ITT in (L) were determined through non-parametric tests. Data in (B-F), (H), and (I-J) were analyzed by one-way ANOVA followed by Bonferroni's multiple-comparison test. Data in (K) and (M) were determined through two-way ANOVA followed by Bonferroni's multiple-comparison test. Bar graphs in AUC of GTT in (L) were calculated using two-sided unpaired t-tests.

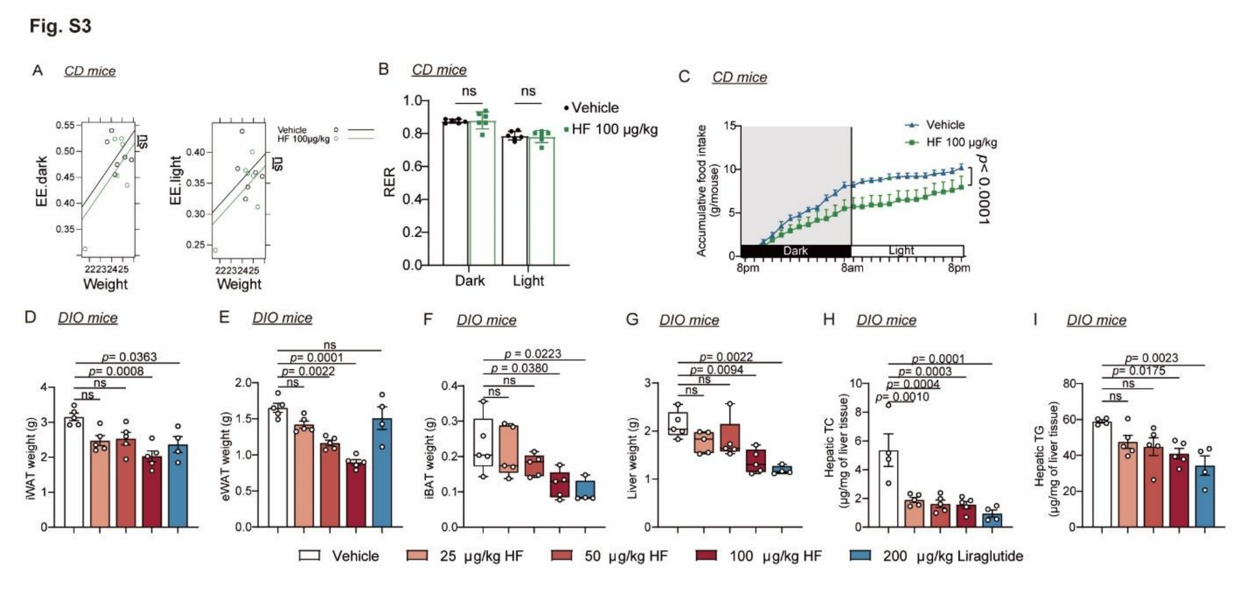


**Fig. S3. Halofuginone ameliorates metabolic dysfunction in mice.**

**(A-C)** 8-week-old male C57BL/6J mice fed chow diet were injected intraperitoneally with vehicle or 100 μg/kg HF in metabolic chamber. **(A)** EE, **(B)** RER, and **(C)** food intake were measured (n=6). **(D-I)** It is the same batch of mice as Fig S2 (A-J). The mice were subjected to the following measurements: **(D)** Weights of iWAT (vehicle or HF group: n = 5, liraglutide group: n = 4). **(E)** Weights of eWAT (vehicle or HF group: n = 5, liraglutide group: n = 4). **(F)** Weights of iBAT (vehicle or HF group: n = 5, liraglutide group: n = 4). **(G)** Liver weights and liver weight normalized by body weight (vehicle or HF group: n = 5, liraglutide group: n = 4). **(H-I)** Hepatic TG and TC (vehicle group: n = 4, HF group: n = 5, liraglutide group: n = 4). Data are presented as mean ± s.e.m. Data in (A) were determined through ANCOVA using body mass as a covariate. Data in (B-C) were determined through two-way ANOVA followed by Bonferroni's multiple-comparison test. Data in (D-E) and (H-I) were analyzed by one-way ANOVA followed by Bonferroni's multiple-comparison test. Data in (F) and (G) were analyzed by non-parametric tests.

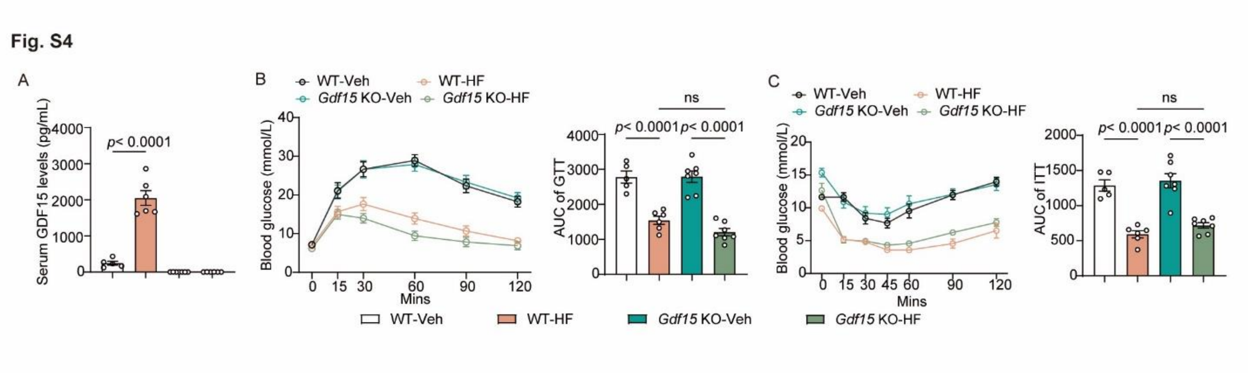


**Fig. S4. Metabolic indices of *Gdf15* KO mice treated with vehicle or halofuginone.**

**(A-C)** 8-week-old male C57BL/6J mice (wild-type) and *Gdf15*$^{-/-}$ mice were fed with HFD for 8 weeks. Wild-type and *Gdf15*$^{-/-}$ mice were then randomly assigned to vehicle or HF (100 μg/kg) group and injected every two days for 8 weeks. **(A)** Serum GDF15 protein levels. **(B)** GTT and **(C)** ITT of mice were detected at weeks 7-8 of administration (WT-Veh group: n = 5, WT-HF group: n=6, Gdf15 KO group: n=7). Data are mean ± s.e.m. *P* values of (A-C) were determined through one-way ANOVA followed by Bonferroni's multiple-comparison test.

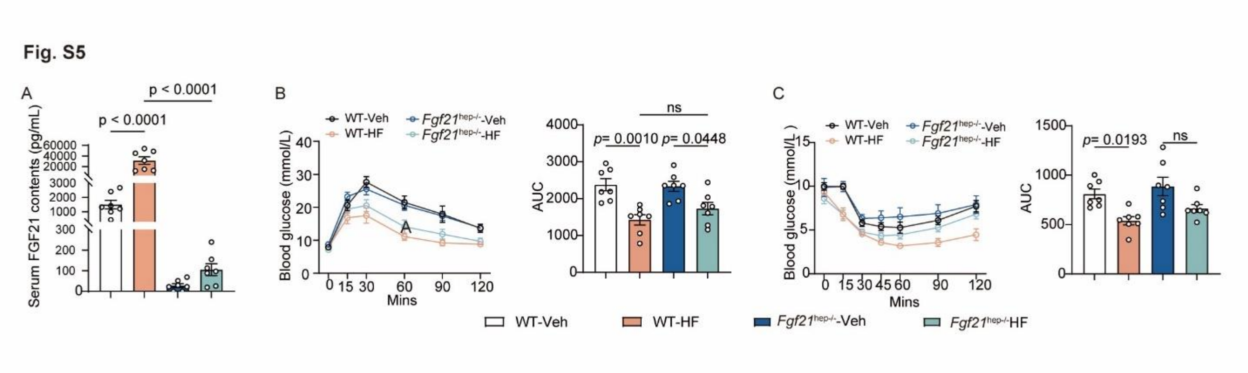


**Fig. S5. Metabolic parameters of WT and *Fgf21*$^{hep-/-}$ mice treated with vehicle or halofuginone.**

**(A-C)** 8-week-old male *Fgf21*$^{flox/flox}$ mice (wild-type) and *Fgf21*$^{hep-/-}$ mice were fed with HFD for 12 weeks, and then randomly divided into vehicle or HF (100 μg/kg) group and injected every two days for 10 weeks. **(A)** Serum FGF21 protein levels. **(B)** GTT and **(C)** ITT were detected at weeks 9-10 of administration (n=7). Data are mean ± s.e.m. *P* values of (A) and (B) were determined through one-way ANOVA followed by Bonferroni's multiple-comparison test.
